# Effect of the divalent cations zinc and calcium on the structure and mechanics of reconstituted vimentin intermediate filaments

**DOI:** 10.1101/844167

**Authors:** Huayin Wu, Yinan Shen, Dianzhuo Wang, Harald Herrmann, Robert D. Goldman, David A. Weitz

**Affiliations:** John A. Paulson School of Engineering and Applied Sciences, Harvard University, Cambridge, MA 02138; Department of Physics, Harvard University, Cambridge, MA 02138; Department of Physics, University of Illinois at Urbana-Champaign, Urbana, IL 61801; Division of Cell Biology, German Cancer Research Center (DKFZ), D-69120 Heidelberg, Germany; Institute of Neuropathology, University Hospital Erlangen, Friedrich-Alexander Universität Erlangen-Nürnberg, Erlangen, Germany; Department of Cell and Developmental Biology, Northwestern University Feinberg School of Medicine, Chicago, IL 60611

**Keywords:** cytoskeleton, rheology, cell mechanics

## Abstract

Divalent cations in a concentration-dependent manner behave as effective crosslinkers of intermediate filaments (IFs) such as vimentin IF (VIF). These interactions have been mostly attributed to their multivalency. However, ion-protein interactions often depend on the ion species, and these effects have not been widely studied in IFs. Here we investigate the effects of two biologically important divalent cations, Zn^2+^ and Ca^2+^, on VIF network structure and mechanics *in vitro*. We find that the network structure is unperturbed at micromolar Zn^2+^ concentrations, but strong bundle formation is observed at a concentration of 100 μM. Microrheological measurements show that network stiffness increases with cation concentration. However, bundling of filaments softens the network. This trend also holds for VIF networks formed in the presence of Ca^2+^, but remarkably, a concentration of Ca^2+^ that is two orders higher is needed to achieve the same effect as with Zn^2+^, which suggests the importance of salt-protein interactions as described by the Hofmeister effect. Furthermore, we find evidence of competitive binding between the two divalent ion species. Hence, specific interactions between VIFs and divalent cations are likely to be an important mechanism by which cells can control their cytoplasmic mechanics.

**Significance:** Intermediate filaments are key structural elements within cells; they are known to form networks that can be crosslinked by divalent cations, but the interactions between the ions and the filaments are not well understood. By measuring the effects that two divalent cations, zinc and calcium, have on the structure and mechanics of vimentin intermediate filaments (VIFs), we show that although both have concentration-dependent effects on VIFs, much more calcium is needed to achieve the same effect as a small amount of zinc. Furthermore, when mixtures of the ions are present, the results suggest that there is binding competition. Thus, cells may use the presence of different cation species to precisely control their internal mechanical properties.

## Introduction

Intermediate filaments (IFs) are one of the major cytoskeletal proteins and are key contributors to the complex internal mechanics of eukaryotic cells (1, 2). Vimentin intermediate filaments (VIFs) are the most abundant IFs in cells of mesenchymal origin, comprising major cytoskeletal systems in numerous cell types, including fibroblasts in connective tissues and endothelial cells lining blood vessels. Vimentin assembles into long polymeric 10 nm filaments that entangle and form complex network structures throughout the cytoplasm. Disruptions in the normal assembly and organization of VIF occur in diseases such as cataracts (3), neuromuscular disorders (4), and some cancers (5). Furthermore, the switching-on of vimentin expression is a marker of cells that are undergoing an epithelial-to-mesenchymal transition, which is accompanied by significant changes in cell stiffness and morphology (6–8). Thus, the structure and mechanics of the VIF networks are essential to their function in cells. These properties of VIFs are thought to be influenced in vivo by post-translational modifications such as phosphorylation. However, because VIFs are highly charged polymers, charge-induced interactions may also play an important role. In fact, purified vimentin protein can be fully assembled from a soluble, tetrameric state that is stably sustained in low ionic strength buffer. Under these conditions VIF networks can be assembled from the tetrameric state *in vitro* solely by tuning the monovalent salt concentration to physiological levels in a pH neutral buffer (9, 10). Thus, the intracellular ionic environment, which is actively regulated by cells, is likely to be critical in determining VIF network properties *in vivo*. Ion pumps in the cell membrane control the concentrations of not only monovalent ions but also some divalent ions such as calcium, whereas other divalent ions such as zinc are balanced using transporter proteins. Due to their multiple charges, multivalent ions are, in contrast to monovalent ions, more capable of mediating interactions between neighboring protein units. For example, millimolar concentrations of Mg^2+^ and Ca^2+^ have been demonstrated to effectively crosslink VIF in their C-terminal domains and this has been shown to stiffen reconstituted VIF networks without changing their structure (11). Further increasing the Mg^2+^ concentration can induce bundling due to counterion-mediated condensation (12–14). There is also evidence that Zn^2+^ may interact with a cysteine in the rod domain, C328, to regulate VIF organization in cells (15). Interestingly, divalent cation species appear to exhibit increasing effectiveness at promoting VIF aggregation according to their atomic number (14). Studies of other proteins have shown that ion-protein interactions can strongly depend on the ion species even when valency is unchanged (16, 17); this is often referred to as the Hofmeister effect. Corresponding species-specific interactions have not been widely explored in VIF networks. Furthermore, the effects of exposing VIFs to multiple types of divalent cations simultaneously, which occurs naturally in the cytoplasm, are also not well known. Characterizing the exact influence that different ions have on VIF networks, both separately and together, is a necessary step toward understanding the mechanical role of IF in cells.

In this work, we investigate the effects of Zn^2+^ and Ca^2+^ cations on VIF network structure and mechanics using VIF networks that have been assembled *in vitro*. We demonstrate that VIFs aggregate and align to form bundles in response to increasing concentrations of Zn^2+^ and Ca^2+^. However much higher concentration of Ca^2+^ compared to Zn^2+^ are required for bundling. Microrheological experiments show that VIF networks stiffen with increasing cation concentrations but once bundles of VIFs form, the networks soften. Furthermore, in mixed ion solutions, we find that the network stiffness depends strongly on the relative amounts of each ion type, suggesting that there is competitive binding between ion species. Interactions between VIFs and divalent cations are likely to be an important mechanism by which cells can control their cytoplasmic mechanics.

## Materials & Methods

### Protein purification and reconstitution

Human vimentin protein is expressed in bacteria (*Escherichia coli*, strain TG1) and purified from inclusion bodies as previously described (18). The protein is stored at -80 °C in 8 M urea, 5 mM Tris-HCl (pH 7.5), 1 mM EDTA, 0.1 mM EGTA, 1 mM DTT, and 10 mM methyl ammonium chloride (MAC). The purity of the protein is verified by SDS–polyacrylamide gel electrophoresis. Before use, the protein is dialyzed into 5 mM Tris-HCl (pH 7.5), 1 mM EDTA, 0.1 mM EGTA, 1 mM DTT in a stepwise manner using membranes with a 20 kDa cut-off. To achieve protein concentrations appropriate for dilution, the protein stock is concentrated using a centrifuge-based filter unit (Ultra-15 Centrifugal Filter Units, Amicon, EMD Millipore, Burlington, MA). The final protein stock concentration is determined by measuring the absorption at 280 nm (Nanodrop ND-1000, ThermoScientific Technologies, Inc., Wilmington, DE).

### Filament assembly and electron microscopy

To improve retention of material on the transmission electron microscopy (TEM) grids and observation of individual filaments, TEM samples were prepared with 0.2 mg/ml vimentin. Assembly was initiated by the addition of a salt buffer with a final Tris-HCl concentration of 25 mM and varying amounts of CaCl2 (Sigma, Burlington, MA) and/or ZnCl2 (Sigma, Burlington, MA). The sample was allowed to polymerize without disturbance at 37 °C for 1.5 hours. A 10 μL sample was applied to a freshly glow-discharged carbon-coated copper or nickel grid and allowed to attach for one minute. The sample was crosslinked by 0.1% glutaraldehyde (Electron Microscopy Sciences, Hatfield, PA) followed by negative staining of the proteins with NanoVan (Nanoprobes, Yaphank, NY). The samples were imaged on a JEOL-2100 TEM operated at 80 keV.

### Image Analysis

Due to the negative EM stain, filaments appear as two dark lines surrounding a brighter core. We obtain the filament widths by plotting the intensity profile of a thin rectangle drawn perpendicular to a filament and taking the width of the peak corresponding to the core. Within the rectangle, the intensity is averaged along the direction of the filament to decrease noise. The analysis is performed using the Fiji distribution of ImageJ (19).

### Microrheology

All mechanical measurements are performed using 1mg/mL vimentin. We coat 3.22 μm fluorescent carboxylated microspheres (Thermo Fisher, Waltham, MA) with PEG using an EDC/NHS reaction to covalently link amine-terminal 2 kDa PEG (Rapp Polymere GmbH, Tuebingen, Germany) to the carboxyl groups on the microsphere surface as previously described (20). We incorporate the PEGylated microspheres into the vimentin assembly buffer before mixing it with the protein stock to the final working concentration. After thoroughly mixing, the solution is immediately transferred to a small well created by a double-sided adhesive silicone spacer (Life Technologies, Waltham, MA) on a No. 1.5 glass cover slip. The sample is sealed using another glass coverslip and allowed to polymerize at 37 °C in the dark. The samples are kept on a slowly rotating platform to prevent settling of the beads. After 1.5 h, the samples are imaged using an epifluorescence microscope (Zeiss Axiovert) outfitted with an LED light source (Colibri2.0) and a low-light camera (Hamamatsu Orca Flash v4). We take images of the fluorescent bead channel every 10 ms for about 20,000 frames. We track the particle positions as well as calculate the MSDs and rheological properties using a customized version of the MATLAB code released by the Kilfoil lab, which is based on the IDL code from Crocker/Weeks (21, 22).

## Results and Discussion

### Zn-VIF networks form bundles at high Zn^2+^ concentrations

To generate VIF networks for structural and mechanical characterization, we use recombinant human vimentin protein that is expressed in *Escherichia coli* (*E. coli*) and purified from inclusion bodies as previously described (18). For reconstitution of functional complexes, the purified protein is dialyzed into a low-salt buffer, and filament assembly is induced by adding a polymerization buffer to a final concentration of 25 mM Tris-HCl (pH 7.5) and 160 mM NaCl. We view the filaments using a transmission electron microscope operated at 80keV to minimize damage to the proteins. Using a standard assembly protocol, the resulting VIF networks form a mesh of smooth, freely organized filaments in the absence of divalent cations. The randomly oriented VIFs are entangled with each other, with many overlapping points and some looped structures, as seen in the electron micrograph in Figure 1A. VIFs are characterized by their length, width, and curvature. With only monovalent cations, VIFs are long and appear straight within a ~1 μm field of view, which is consistent with previous measurements that find the persistence length of vimentin to be around 1μm (10). To determine the widths of VIFs, we plot the intensity profile perpendicular to individual filaments at multiple locations along its length and measure the width of the bright peak that corresponds to the filament core. We find that there is some variation in the diameter along individual filaments and that they are 10.4 +/- 1.6 nm wide on average (Figure 1B). This average width is fully consistent with previous observations of VIFs (9, 10).

**Figure 1.**
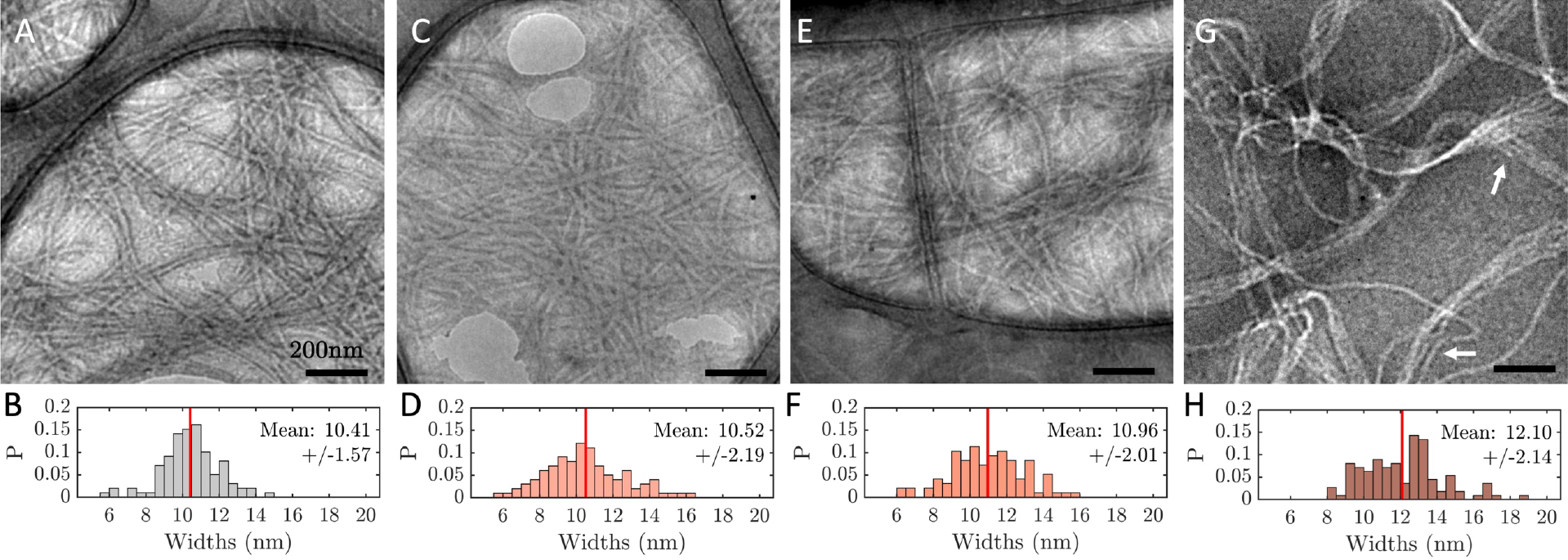
Electron micrographs of VIF with various molar ratios of Zn^2+^ and the corresponding filament width distributions: a-b) *R*=0, c-d) 0.054, e-f) 0.54, g-h) 5.4. At the highest concentration of Zn^2+^, the average filament width increases. The scale bars represent 200 nm. Arrows indicate examples of distinct bundles.

When ZnCl2 is included in the polymerization buffer at a cation-to-vimentin molar ratio of *R_Zn_* = 0.054, there are no significant changes in the overall network structure (Figure 1C) and the size of individual VIFs (Figure 1D). Similarly, at *R_Zn_* = 0.54, the polymerized VIF network structure remains similar (Figure 1E) although individual filaments widen slightly (Figure 1F).

Upon increasing the Zn^2+^ concentration to *R_Zn_* = 5.4, there is a marked difference in the network structure: the empty space between VIFs becomes much larger and the most prevalent structure is no longer comprised of individual ~10 nm filaments. Instead, VIFs are aligned and closely packed in bundles over micron-long distances, as seen in Figure 1G. Many of the bundles appear like flat sheets, although this could be an artifact of the staining and drying process. The bundles twist and cross over each other, as highlighted by the upper arrow in Figure 1G. The bundle structures are heterogeneous, containing as many as 5 VIFs each. Furthermore, the number of VIFs varies along the entire length of a single bundle and individual VIFs can be part of multiple distinct bundles at different locations (see lower arrow in Figure 1G). The morphology of the VIF bundles is consistent with reports that multivalent ions can cause neighboring filaments to aggregate through coordinating lateral interactions of filaments along their lengths, which causes them to line up in a parallel manner (12, 23). Due to the lateral alignment, individual filaments are well-defined even when they are in bundles, and thus the dimensions of individual filaments can be determined. We find that the filaments widen from 10.4 +/- 1.6 nm to 12.1 +/- 2.1 nm (Figure 1H). However, the filaments remain very long, suggesting that the bundling process does not interfere with the elongation step of filament assembly.

### Networks crosslinked by Zn2+ are stiff but VIF bundles are soft

We use microscale measurements to characterize the impact of Zn^2+^ ions on VIF network mechanics. The network properties can be probed by observing the thermally-driven motion of tracer particles within the network (24, 25). To do this, we incorporate 3.22 μm PEG-functionalized particles into the vimentin solutions as they polymerize and then capture videos of the particle positions over the course of several minutes. The surface layer of PEG prevents the particles from interacting with vimentin, which could influence the structure as the network forms. Assuming a homogeneous network made from 1mg/mL of vimentin, the expected mesh size, ~ 0.45 μm (11), is much smaller than the particle diameter, ensuring that the particles are fully trapped by the network. Based on the electron microscopy results, this assumption is likely to hold for *R_Zn_* < 5.4, but may not apply to bundled VIFs.

To quantitatively understand the particle motion, we calculate the mean-squared displacement (MSD) for each particle. Individual and average MSDs for a VIF network without divalent ions are plotted as a function of lag time *τ* in Figure S1. There is not much variation between individual particle MSDs, indicating that the network is fairly homogeneous. The MSDs are subdiffusive and approach a plateau at larger values of *τ*, indicating that the particles are being influenced by the network. Embedded particles are only able to move as much as thermal energy can deform the network. Therefore, the network elasticity determines the amount of confinement. Since there are no crosslinks, this confinement likely arises from entanglements within the network, which have been shown to contribute significant elasticity to other networks of semiflexible polymers such as actin filaments (25). By plotting the particle location over time, which shows its trajectory, we find that the path covers a circular area, without specific directionality, and that it is smaller than the particle size. The trajectories of two different tracers are shown in Figure 2A.

**Figure 2.**
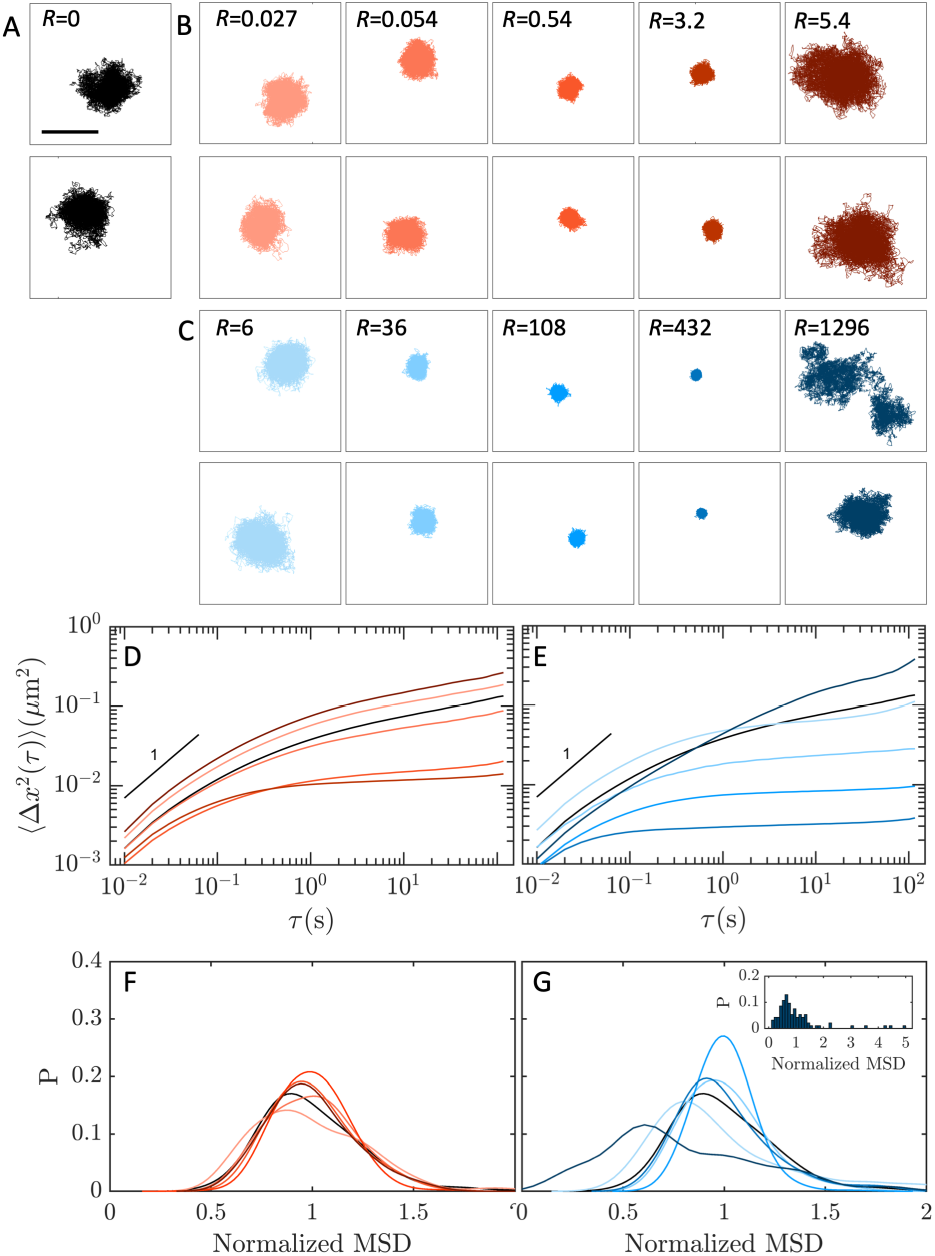
Sample particle trajectories for two particles in A) VIFs without divalent cations, B) VIFs with Zn^2+^, and C) VIFs with Ca^2+^. The scale bar represents 1μm and the molar ratios are as indicated. Average MSD curves for Zn-VIF and Ca-VIF networks are shown in D and E, respectively; the color of each curve corresponds to the divalent cation concentration, increasing from lightest (*R_Zn_* = 0.027, *R_Ca_* = 6) to darkest (*R_Zn_* = 5.4, *R_Ca_* = 1296). *R* = 0 is plotted in black as a reference. Normalized distributions of the MSDs at *τ* = 22 s are shown using the same color scale in F and G. The distribution has a long tail to the right at *R_Ca_* = 1296, shown in the inset of G.

When *R_Zn_* < 5.4, the particle motion progressively covers less area with increasing Zn^2+^ concentration (Figure 2B). Since the network structure does not change within this concentration range, this trend reflects changes in the mechanical properties. We find that the average MSDs of Zn-VIF networks have more pronounced plateaus as *R_Zn_* increases between 0.027 and 3.2 (Figure 2D). This suggests that the particles are caged by an increasingly elastic network rather than slowed down by a more viscous-like environment. Thus, changes in cation concentration significantly affect the local environment experienced by the tracer particles, even at concentrations that are too low to alter the network structure. After the VIF network bundles, however, the particle motion increases again (Figure 2B). The MSD curve also approaches but does not fully reach a plateau, similar to the curve obtained for a VIF network without any divalent cations (Figure 2D). These effects may reflect an increased mesh size, a weaker network, or a combination of both; since we do not observe any particles freely diffusing between microenvironments, we conclude that the mesh size here remains smaller than the particle size.

Based on particle-to-particle variations of the MSDs of individual particles, we can also gain insight into the network heterogeneity. For each particle in a Zn-VIF sample, we take the MSD at *τ* = 22s and normalize it by the sample average. For all samples, we find that the distributions of normalized MSDs are nearly symmetric around 1 and have similar widths, as seen in Figure 2F. This is true even for the bundled network, indicating that the networks are homogeneous on the scale of the probe particles.

We measure the network rheological properties from the particle displacements. The MSDs of individual particles may be strongly influenced by a number of factors on the scale of the particle size, including variations in particle size or shape, and local heterogeneities in network structure. For this reason, we use two-point microrheology, which considers only the correlated motion of particles within a sample to probe the material over length scales much larger than the particle size (24). This method is therefore not dependent on the probe size or network heterogeneities and is able to reflect the bulk mechanical properties. We calculate a two-point MSD for each sample and use the generalized Stokes-Einstein relation (GSER) to determine the network storage modulus, *G′*(*ω*), and loss modulus, *G″*(*ω*), which represent the elastic and viscous properties, respectively(26). Without divalent cations, the VIF network is predominantly elastic, as evidenced by a dominant *G′* at intermediate frequencies. The network stiffness is slightly frequency-dependent, varying as *G′* ~ *ω*^0.2^, which suggests that the VIF network consists of VIFs that are entangled but not crosslinked. It is also very soft, *G′* = 0.009 Pa at *ω* = 1 rad/s, as shown in Figure 3A (black line). This value is in good agreement with semiflexible polymer theory, which predicts a stiffness of ~0.01 Pa based on the persistence length of vimentin, *l_p_* ~ 0.5-1 μm (27, 28). We note here that this network stiffness is much lower than previously reported values measured by bulk rheometers (29, 30). There is some evidence that bulk measurements of very soft gels that have *G′* < 1 Pa may over report *G′* due to elastic contributions by the interface at the exposed sample edge (31).

**Figure 3.**
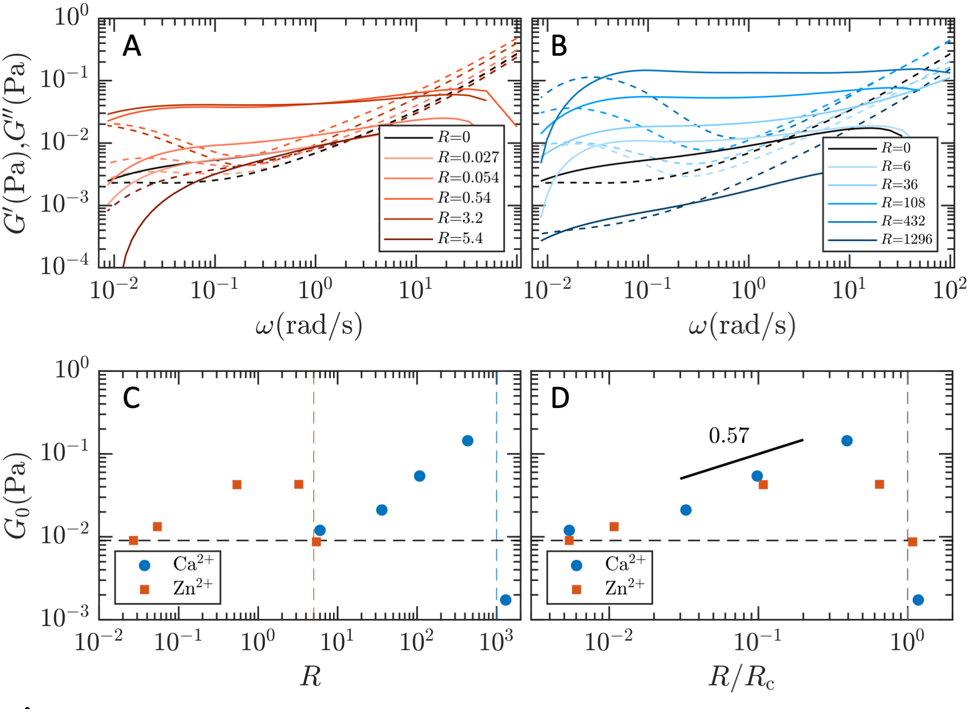
Rheological properties G′ and G″ of the VIF networks are plotted as a function of frequency ω. Zn-VIF (A) and Ca-VIF (B) networks both behave as crosslinked networks, displaying decreased frequency dependence and increased stiffness. C) The plateau modulus *G_0_* varies with molar ratio *R* in a similar way for both Zn^2+^ and Ca^2+^, first increasing then dropping suddenly. D) The data can be collapsed when *R* is scaled by a bundling concentration, *R_c_*, for each ion species.

Zinc cations strongly affect the mechanical properties of VIF networks. Below a molar ratio of 5.4, the VIF networks stiffen and become less frequency-dependent as *R_Zn_* is increased (Figure 3A), which suggests that Zn^2+^ ions effectively behave as crosslinkers of VIFs. Since the network structure does not change within this range, the differences in *G′* likely reflect an increased crosslinking density. However, when *R_Zn_* reaches 5.4, we find that the network softens dramatically such that it is similar to an entangled VIF network. The two-particle analysis considers only the portion of particle motion that is due to network fluctuations and is therefore not influenced by any uncaging events. Thus, the lower value of *G′* reflects changes in the network itself, including an altered mesh size or VIF properties. The bundled Zn-VIF network is more frequency-dependent than the entangled one, falling off faster at low frequencies, and it is barely elastic within a small frequency range, likely due to the coarser network structure.

The values for *G′* and *G″* cross at low frequencies due to relaxation along the filaments through network constraints. The crosslinked networks appear to relax somewhat more quickly than entangled VIFs, which may reflect the transience of divalent ion crosslinks (32). However, the total time of the measurements does not offer enough statistics to establish any significant difference between the samples; all of the networks relax very slowly, on the order of 1-2 minutes, whether crosslinked or not.

### Ca^2+^ affects VIFs in a similar manner as Zn^2+^ but at higher concentrations

There is some evidence that Ca^2+^ also crosslinks vimentin (29, 33), but its effect has not been fully characterized, particularly at these concentrations. Thus, we incorporate CaCl2 into the VIF polymerization buffer to determine its influence on network structure and mechanics. By using the chloride salt, we preserve the counterion identity, which facilitates direct comparisons with the effects of ZnCl_2_. Electron micrographs of VIF networks assembled at high molar ratios of Ca^2+^ show large bundles, thickened filaments, and a sparser mesh (Figure S2). These results are qualitatively similar to the Zn-VIF bundles but the concentration of Ca^2+^ used here (*R_Ca_* ~ 2000) is much larger, by over 2.5 orders of magnitude, than the concentration of Zn^2+^ at which visible bundling occurs (*R_Zn_* = 5.4).

Particles embedded within Ca-VIF networks of varying *R_Ca_* again reveal similar behavior as those in Zn-VIF networks, but at very different cation concentrations. As the Ca^2+^ concentration increases from *R_Ca_* = 6 to 432, particle motion decreases significantly (Figure 2C). Interestingly, the degree of confinement is greater at *R_Ca_* = 432 than in any of the Zn-VIF networks. The corresponding MSDs all show plateaus that reflect the constraint of particles by the surrounding network (Figure 2E). At *R_Ca_* = 1296, however, particle motion increases again and some particles can permeate through the network (Figure 2C), suggesting that the network has formed bundles and the mesh size is now close to the bead size. The shape of the ensemble-averaged MSD also changes at this molar ratio; it remains slightly subdiffusive for all lag times but continues to increase over time (Figure 2E). Particles in this network exhibit a broader distribution of MSDs compared with those embedded within networks at lower *R_Ca_*, which suggests increased heterogeneity due to bundling (Figure 2G); furthermore, the few highly motile particles cause the peak to shift left, and the long tail can be seen in the inset of Figure 2G.

The rheology of Ca-VIF networks is also strongly dependent on the divalent cation concentration. VIFs assembled in the presence of Ca^2+^ generally form elastic networks that are frequency-independent and stiffer than entangled VIFs, reminiscent of crosslinked gels. However, the bundled Ca-VIF, like its Zn-VIF counterpart, forms the softest network, as seen in Figure 3B.

Since *G′* exhibits little frequency-dependence at intermediate frequencies, we have determined a plateau elastic modulus, *G_0_*, at *ω* = 1 rad/s. In the range 0.027 < *R_Zn_* < 0.54, we find that *R_Zn_* directly influences *G_0_* according to the relation *G_0_* ~ *R*_*Zn*_^0.51^. Similarly, we find that Ca^2+^ stiffens the networks according to *G_0_* ~ *R_Ca_*^*0*.58^ in the range 36 < *R_Ca_* < 432, which agrees with bulk rheology studies on Ca^2+^ and Mg^2+^ (29, 33). Remarkably, although the exponents are similar for Zn^2+^ and Ca^2+^, their absolute concentrations differ by more than two orders of magnitude (Figure 3C). In fact, Zn^2+^ induces VIF bundling at molar ratios where Ca^2+^ is only beginning to stiffen the network.

When the network is formed with Zn^2+^, *G_0_* appears to saturate between the *R_Zn_* = 0.54 and 3.2. By contrast, the Ca-VIF networks do not exhibit a plateau at higher *R_Ca_* but instead stiffen monotonically. Moreover, calcium stiffens the VIF network more than zinc does. These differences in stiffening behavior suggest subtle differences in the way each ion species binds to vimentin. VIF structure reflects a balance between the processes of lateral assembly and filament elongation, which may be influenced by both the valency and the concentration of the counterions. Here, we use a physiological concentration of monovalent ions and add several orders of magnitude fewer divalent cations. Under these conditions, divalent cations are presumed to primarily serve in a crosslinking capacity with minimal effects on VIF structure (29, 34). However, it is likely that the divalent cations facilitate crosslinks or lateral interactions between vimentin tetramers even as the filaments are assembling (9, 34). Such effects will affect the network mechanics in a way that depends on the comparative strengths of interaction as well as the number of interactions. Thus, although Zn^2+^ binds very strongly to vimentin, its effect on VIF network mechanics may plateau once the network branch points are saturated with crosslinks since the numbers of divalent ions and vimentin monomers are comparable. In the case of Ca^2+^, the large number of divalent ions can coordinate to stiffen the network even more.

For both ion species, a network of bundles is softer than the crosslinked networks. Since the total amount of protein is unchanged, the network of bundles must have a larger mesh size. The plateau elastic modulus of a crosslinked semiflexible polymer network can be described by 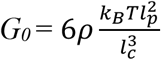 where *l_c_* is the crosslinking length and *ρ* is the total length of filaments per unit volume (27). In a densely crosslinked network, *l_c_* is the same as the entanglement length *l_e_*, which scales as *l_p_*^1/5^*ρ*^-2/5^. As bundles are introduced, the network structure is now defined by the bundles rather than by the individual filaments; thus, the new total length is *ρ_bundle_* = *ρ/N* where *N* is the number of filaments per bundle, and *l_p_* scales as *N^x^*, where *x* represents the inter-filament coupling strength. The exponent ranges between 1 and 2 for loosely- and tightly-coupled bundles, respectively (35). Therefore, the plateau elastic modulus scales according to 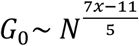, which predicts network softening if bundles are loose (*x* = 1) and stiffening if the coupling is tight (*x* = 2). Since the *G_0_* of VIF networks decreases upon bundling with both cation species, we conclude that the ionically-bundled filaments are loosely coupled.

Overall, the shape of the *G_0_* vs *R* curves are similar for both zinc- and calcium-modified networks. We find that the points collapse and follow an exponent of about 0.57 when the molar ratios are rescaled by a threshold molar ratio, *R_c_* (Figure 3D). This suggests that the mechanism of stiffening may be fundamentally similar but that different interaction strengths could determine the relevant concentration range. From the rescaled data, we estimate the species-specific bundling thresholds to be approximately *R_Zn_* = 5 and *R_Ca_* = 1000.

### Ionic properties affect interactions within VIF networks

The vast difference in the crosslinking and bundling concentrations suggest that Zn^2+^ and Ca^2+^ interactions with vimentin are not purely electrostatic in nature. This result is reminiscent of the Hofmeister series, in which ion species are ordered according to their ability to interact with proteins, a property that still cannot be explained from first principles (17, 36). The Hofmeister effect is thought to be a complex balance of the many potential interactions in the system, including ion-counterion, ion-solvent, and ion-protein interactions, which necessarily depend on intrinsic properties of the ion beyond just its charge. To explore this further, we consider the effects of the solvent. A typical cytoplasm is mostly water, and the studies above are all performed in a H_2_O-based buffer. However, by exchanging a fraction of H2O for D_2_O, the interactions can be influenced. For example, ion hydration is likely to decrease due to stronger hydrogen bonding in D_2_O.

Therefore, to determine whether an entangled VIF network is altered by the addition of D_2_O, we make microrheological measurements on pure VIF networks polymerized in a buffer that contains 45% D_2_O (Fig 4A). The results show that the network stiffness doubles compared with a network polymerized in H2O, suggesting that there could be strengthening of the protein structure itself due to altered hydrogen bonding within the protein structure. Furthermore, the decreased ability of monovalent ions in D_2_O to induce long-range ordering of water molecules (37) may lead to slight differences in protein assembly that affect the filament properties.

When zinc or calcium divalent cations are introduced into the system, the dependence of stiffness on the ion concentration appears to disappear in both cases; instead, the networks made with each ion species are all about the same stiffness. Notably, this stiffness is slightly higher in the Ca-VIF networks than in the ones prepared with Zn^2+^ (Fig 4B, C), which is reminiscent of the different maximum stiffnesses achieved by each species in a standard buffer. This provides further evidence that calcium ions are able to stiffen VIFs more but that zinc ions are highly effective at very low concentrations. At high *R*, the Ca-VIF network softens, likely due to bundling. Conversely, the Zn-VIF network remains stiff at high *R*, suggesting that D_2_O may stabilize the system against counterion-induced aggregation in some cases. Despite its chemical similarity with H_2_O, D_2_O has significant effects on vimentin subunits, VIFs, and their interactions with ions. Whether these are effects of ion solubility or other ionic characteristics cannot be inferred at this time. Nevertheless, these results provide crucial support for the importance of hydrogen bonding and ion hydration, and by extension the Hofmeister effect, in the interactions between VIF and divalent cations.

### Competitive binding of Zn^2+^ and Ca^2+^

In cells, divalent ion species rarely exist in isolation; instead, the co-existence of many species is much more common. Thus, we study the network mechanics in the presence of mixed ion solutions. Holding the vimentin protein concentration constant, we vary the mixtures of Ca^2+^ and Zn^2+^ using two different concentrations of each ion within the crosslinking regime (*R_Ca_* = 36 and 108, *R_Zn_* = 0.054 and 0.54) and measure the microrheological properties of the resultant networks. To determine the influence of each component, we calculate the fractional change in network stiffness relative to the single-species VIF networks, (*G_0,mix_ - G_0,single_*) / *G_0,single_*. For each combination, we compare *G_0,mix_* with the stiffness of its components, *G_0,Zn_* and *G_0,Ca_*. Naively, we might expect the effects to be additive, as the total divalent ion concentration increases. However, we find instead that the stiffness depends on the concentrations. When *R_Ca_* = 36, the addition of either amount of Zn^2+^ changes the network stiffness by almost 50% compared to the reference Ca-VIF stiffness, as seen in Figure 5A. By comparison, adding Ca^2+^ at a concentration of *R_Ca_* = 36 to Zn-VIF networks results in smaller changes to the network stiffness (Figure 5B). The mixed ion network stiffness is in between those of the single-species Ca-VIF and Zn-VIF networks, but is closer to the Zn-VIF stiffness in both cases, suggesting that vimentin may slightly prefer to bind Zn^2+^ at these concentrations. However, at *R_Ca_* = 108, the mixed ion networks are about the same stiffness as the reference Ca-VIF (Figure 5A), indicating that networks with high concentrations of Ca^2+^ are not sensitive to the presence of Zn^2+^. Correspondingly, when this amount of Ca^2+^ is added to Zn-VIF networks, it results in significant stiffening when *R_Zn_* = 0.054 and slight stiffening when *R_Zn_* = 0.54 (Figure 5B). Thus, it appears that there is competition between the ion species in two regimes: one at lower *R_Ca_* where the single-species network stiffnesses are comparable and Zn^2+^ has a stronger influence, and another at higher *R_Ca_*, where the effect of Ca^2+^ dominates. This result is likely related to the differences in their crosslinking behavior as well as the number of ions present, since Ca^2+^ ions outnumber Zn^2+^ ions by tens to thousands of times.

**Figure 4.**
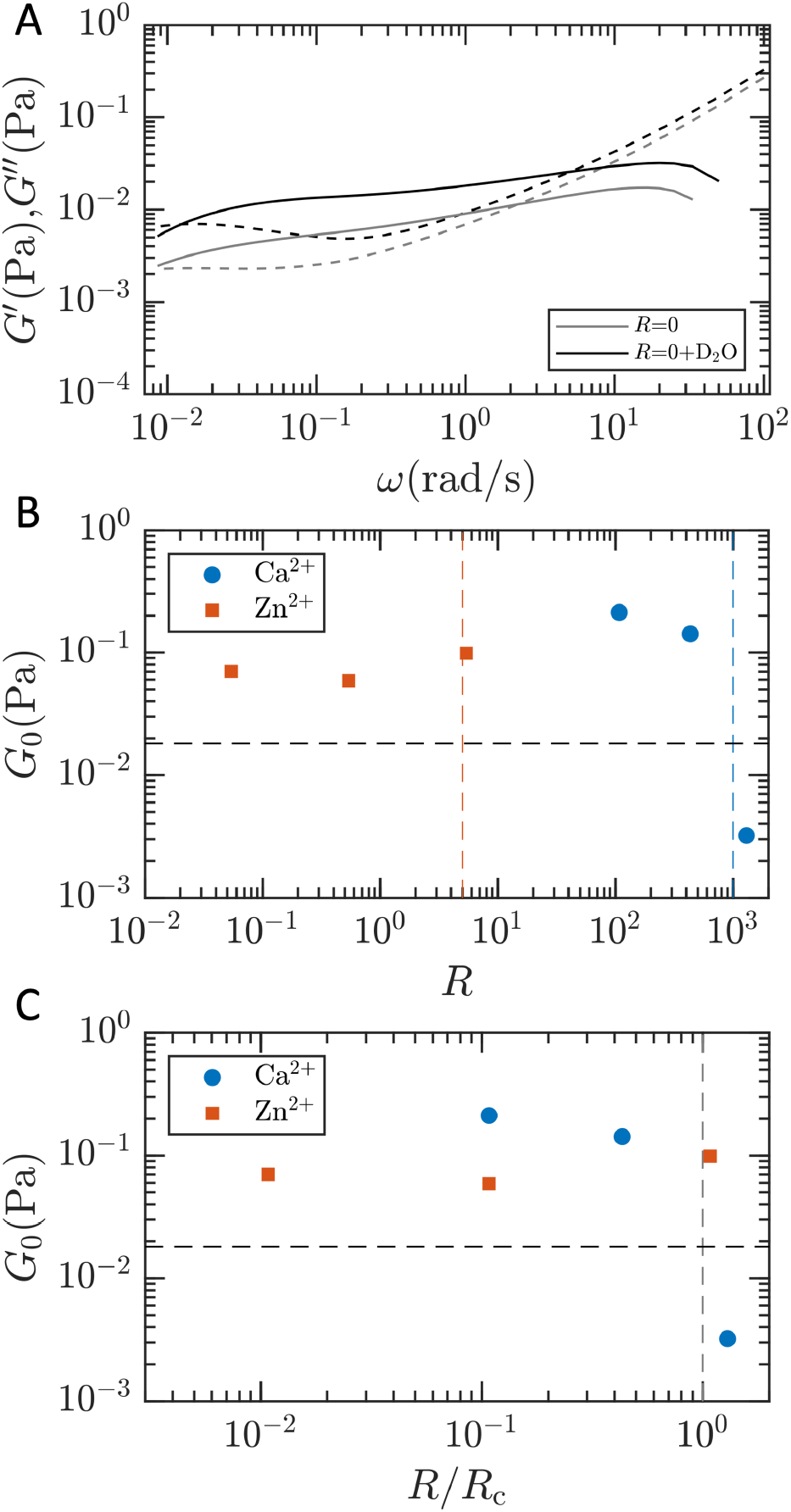
The effects of changing the solvent to 45% D_2_O. A) The VIF network without divalent cations is stiffer. B) *G_0_* is no longer concentration-dependent and networks are stiffer overall. C) Ca-VIF networks are stiffer than Zn-VIF networks.

**Figure 5.**
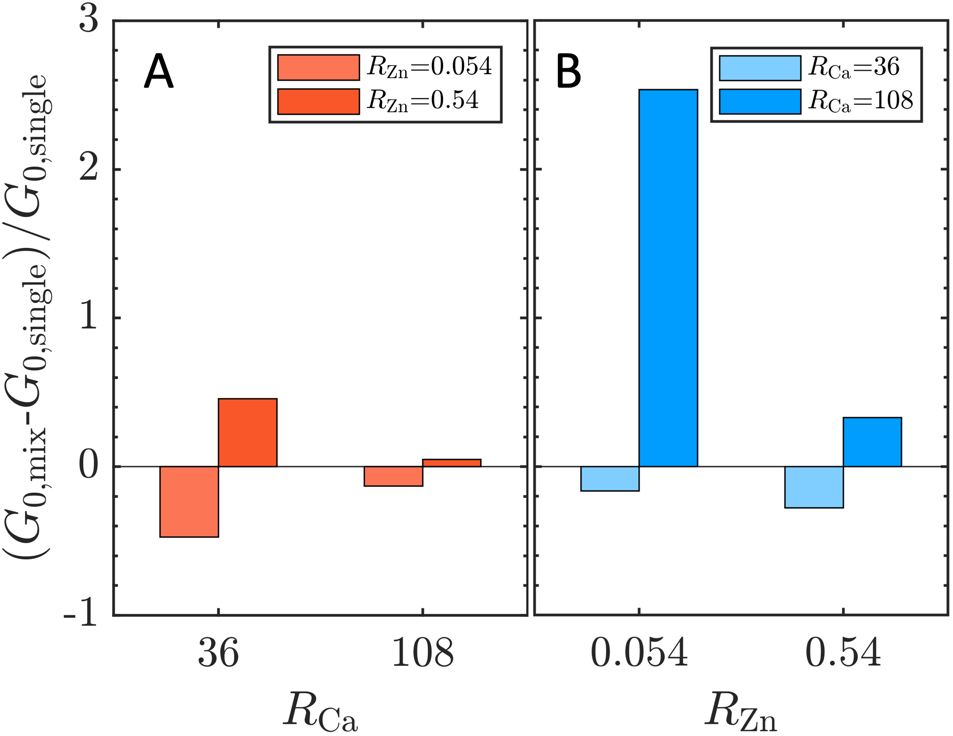
Mixtures of Zn^2+^and Ca^2+^ show binding competition between the ion species. The fractional change in modulus when the mixtures are viewed as A) Zn^2+^ added to Ca-VIF networks, and B) Ca^2+^ added to Zn-VIF networks.

Molar ratios of *R* = 36 and 108 correspond to absolute calcium concentrations of 0.67 mM and 2 mM, respectively. The two Zn^2+^ concentrations are equivalent to 1 μM and 10 μM. By comparison, intracellular concentrations of Ca^2+^ range from 100 nM in cells at rest to millimolar concentrations during cell signaling events or in cases of cell damage (38, 39), whereas the cells typically contain a total of 200-300 μM of Zn^2+^ (40). However, since both Ca^2+^ and Zn^2+^ have important intracellular signaling functions, these concentrations are not uniform throughout the cytoplasm. Instead, there are many mechanisms to bind or sequester the ions, resulting in local concentrations that can be much higher or lower. By preferentially binding different ion species at different concentrations, vimentin subunits and VIFs could also serve in a regulatory or buffering role. As it interacts with the ions, the mechanical properties of the VIF network also change. For example, during a Ca^2+^ influx, the excess Ca^2+^ may further stiffen the VIF network, thereby enhancing protection of the cell against the external stress that triggered the spike. Additionally, the stiffening of VIFs has the potential to affect force propagation through the network as well as its mechanical interactions with other cytoskeletal proteins such as actin. These results point to structural and mechanical adjustments of vimentin as a means by which cells can modify their internal mechanics in response to stress.

## Conclusions

We have shown that the interactions between divalent cations and VIFs do affect the protein structure and mechanics in profound ways. Vimentin is a highly charged polyelectrolyte that forms very flexible polymers. Its flexibility is necessary in cellular functions that involve localizing cytoplasmic organelles and proteins (41, 42), but its mechanical properties are also essential for protecting the cell from external stresses. Here, we see that divalent cations, which are ubiquitous in cells, offer a simple way for the cell to regulate their cytoplasmic properties. By carefully controlling the intracellular ionic environment, cells can tune the VIF network stiffness without changing the network structure, or they can induce bundles that significantly change the structure but are less stiff overall. Furthermore, subtle differences at ion concentrations below the bundling threshold suggest that VIF could act as a zinc-specific buffer, which may be important in some signaling events; this may be due to specific interactions between zinc and the single cysteine residue in the rod domain. Ion-specific effects may also become more important as cells experience forces on different time scales or as deformations become larger. Moreover, as vimentin is not the only filamentous protein in cells but is well integrated into the cytoskeleton, its mechanical properties are bound to affect the entire system via its interactions with the other mechanical elements.

## Author Contributions

H.W. and Y.S. designed and performed experiments, H.W. and D.W. contributed analytic tools, and H.W. analyzed the data. H.W., H.H., R.D.G., and D.A.W. wrote the manuscript.

## Acknowledgments

We thank P. Janmey and S. Köster for helpful discussions. We also thank S. Stoilova-McPhie, C. Marks, and D. Bell for their help with imaging. R.D.G. and D.A.W. were supported by NIH Grant 2P01GM096971-06. DAW was also supported by NSF grant DMR-1708729. H.H. was supported by the German Research Foundation (DFG, HE-1853/11-1). This work was performed in part at the Harvard University Center for Nanoscale Systems (CNS), a member of the National Nanotechnology Coordinated Infrastructure Network (NNCI), which is supported by the National Science Foundation under NSF award no. 1541959.

