## Supplemental Figure S1 for "Effect of the divalent cations zinc and calcium on the structure and mechanics of reconstituted vimentin intermediate filaments"

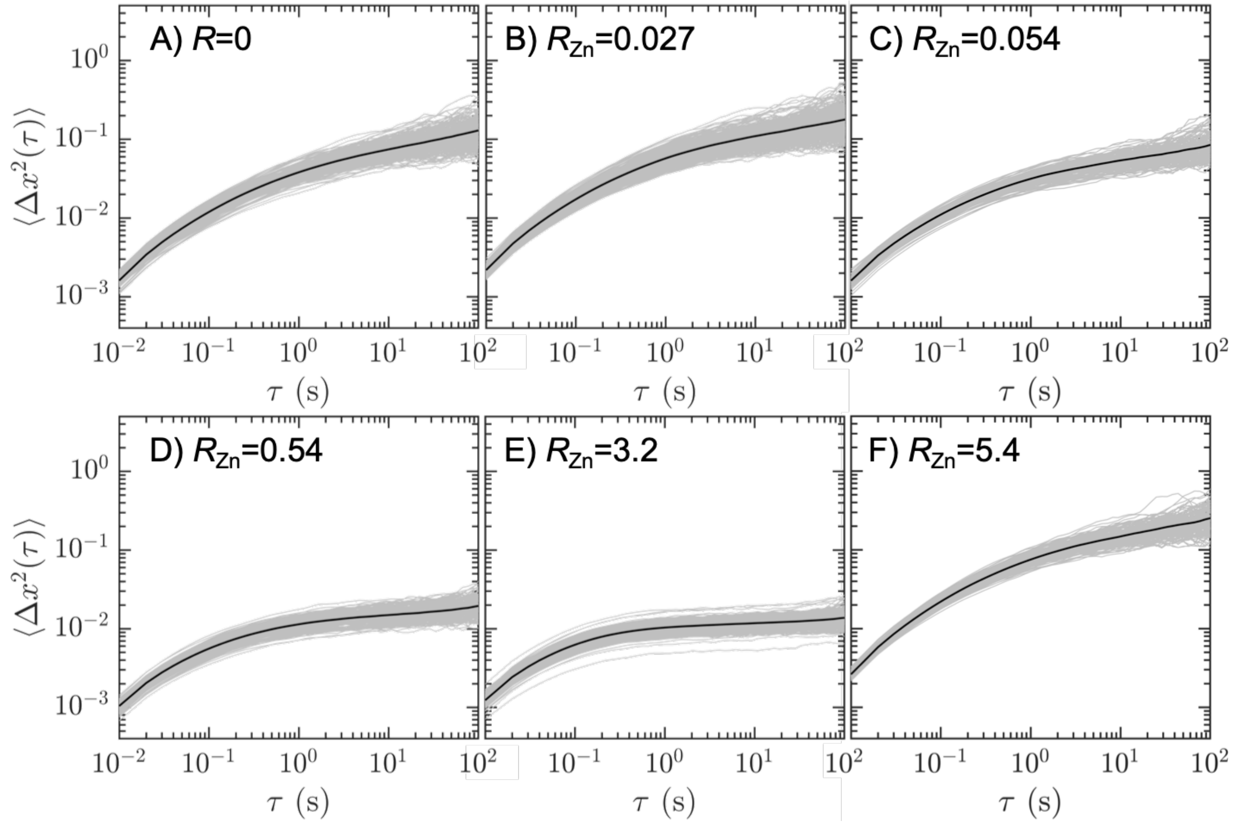


Figure S1.
MSDs for all particles in Zn-VIF networks at molar ratios of A) 0, B) 0.027, C) 0.054, D) 0.54, E) 3.2, and F) 5.4. The black line represents the sample average.
