## Supplemental Figure S2 for "Effect of the divalent cations zinc and calcium on the structure and mechanics of reconstituted vimentin intermediate filaments"

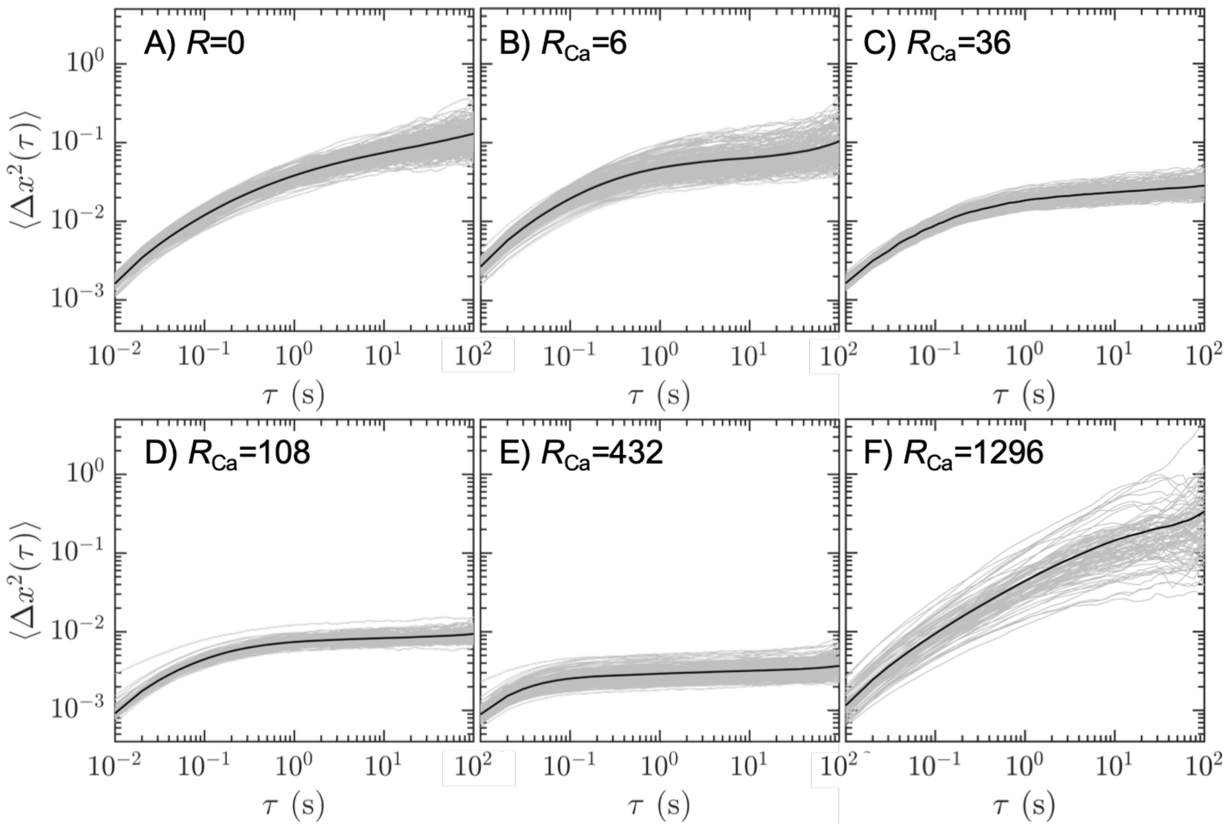


Figure S2.
MSDs for all particles in Ca-VIF networks at molar ratios of A) 0, B) 6, C) 36, D) 108, E) 432, and F) 1296. The black line represents the sample average.
