## Supplemental Figure S3 for "Effect of the divalent cations zinc and calcium on the structure and mechanics of reconstituted vimentin intermediate filaments"

$R_{Ca}$=~2000

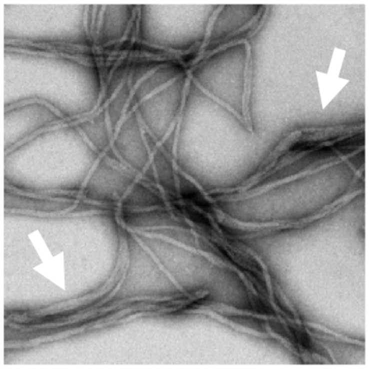

Figure S3.
Electron micrograph of VIFs bundled by $Ca^{2+}$ at a molar ratio of about 2000. Arrows indicate bundles and thickened filaments.
